# LPS binding caspase activation and recruitment domains (CARDs) are bipartite lipid binding modules

**DOI:** 10.1101/2024.10.07.617105

**Authors:** Anh B. Cao, Pascal Devant, Chengliang Wang, Mengyu Sun, Stephanie N. Kennedy, Jianbin Ruan, Jonathan C. Kagan

## Abstract

Caspase-11 is an innate immune pattern recognition receptor (PRR) that detects cytosolic bacterial lipopolysaccharides (LPS) through its caspase activation and recruitment domain (CARD), triggering inflammatory cell death known as pyroptosis. Caspase-11 also detects eukaryotic (*i.e.* self) lipids. This observation raises the question of whether common or distinct mechanisms govern the interactions with self and nonself lipids. In this study, using biochemical, computational, and cell-based assays, we report that the caspase-11 CARD functions as a bipartite lipid-binding module. Distinct regions within the CARD bind to phosphate groups and long acyl chains of self and nonself lipids. Self-lipid binding capability is conserved across numerous caspase-11 homologs and orthologs. The symmetry in self and nonself lipid detection mechanisms enabled us to engineer an LPS-binding domain *de novo*, using an ancestral CARD-like domain present in the fish *Amphilophus citrinellus*. These findings offer critical insights into the molecular basis of LPS recognition by caspase-11 and highlight the fundamental and likely inseparable relationship between self and nonself discrimination.

## Introduction

The accuracy of self-nonself discrimination is central to organismal identity and host defense. Within mammals, these two needs are not distinct, as the accurate identification of self molecules is key for immune system functions in host defense. This “self-first” paradigm was uncovered through studies of how antigen receptor interactions impact lymphocyte development (Koch and Radtke 2011). In the thymus, developing T cells are first selected for the expression of α/β T cell receptors (TCRs) that recognize self-peptides associated with major histocompatibility complex (MHC) (Klein et al. 2014; Koch and Radtke 2011). T cells bearing self-reactive TCRs are subsequently subjected to negative selection to eliminate cells that react excessively to self-components. Only after this process is complete do T cells exit the thymus to survey the body for nonself antigens (Klein et al. 2014; Koch and Radtke 2011). The necessity of self-recognition extends beyond a developmental requirement. The degree of self-reactivity positively correlates with the binding strength of TCRs to foreign antigens (Mandl et al. 2013).

In contrast to the self-first paradigm of lymphocyte functions, the functions of myeloid cells are considered to operate via the opposite principle to govern host defense. Rather than detecting self molecules, macrophages and dendritic cells, for example, use pattern recognition receptors (PRRs) to identify nonself molecules such as microbial cell wall components or nucleic acids (Li and Wu 2021). These PRR ligands are referred to as pathogen associated molecular patterns (PAMPs). The germline encoded nature of genes encoding PRRs has led to the belief that PRRs are fixed in their specificity, whereas each organism has a unique antigen receptor repertoire (Li and Wu 2021). However, PRRs are among of the most rapidly evolving proteins in nature, suggesting that PRR specificity remains subject to evolutionary pressures (G. Liu et al. 2020; Ngou et al. 2024). Such pressures are commonly discussed in the context of infection, whereby PRRs need to evolve to maintain the ability to detect pathogens that have higher rates of mutation and faster replication cycles (G. Liu et al. 2020; Ngou et al. 2024).

The above-described selective pressure relates to the need for a PRR to detect nonself, in order to ensure efficient pathogen detection. Another pressure on PRRs could be the need to avoid the detection of self, in order to prevent pathological inflammatory responses in the absence of infection. In this view, self ligand detection would be an activity that is counter selected for PRRs, but positively selected for in antigen receptors. Yet, instances of self-ligand detection by PRRs are increasingly common. For example, while nucleic acids are not uniquely nonself molecules, DNA and RNA structures are the most common PAMPs in humans and mice, yet (Li and Wu 2021). Nucleic acid sensing PRRs include 5 of the 12 members of the murine Toll-like Receptor (TLR) family, all 3 members of the RIG-I like Receptor family, and the PRRs protein kinase R (PKR), absent in melanoma 2 (AIM2) and cyclic GMP-AMP synthase (cGAS) (Kong et al. 2023; Motwani, Pesiridis, and Fitzgerald 2019; Schlee and Hartmann 2016). As there is no feature of microbial nucleic acids that provides a definitive distinction from self-nucleic acids (Hu et al. 2016; Huang et al. 2020; Kong et al. 2023; Rathinam et al. 2010; Uggenti, Lepelley, and Crow 2019), it is puzzling why the most common PAMPs offer such a blurry line between self and nonself. Stated differently, if self detection activities are counter-selected for during PRR evolution, then nucleic acids would not have evolved as the most common type of PAMP.

The best justification that self-ligand detection is a counter-selected trait relates to PRRs that detect cell wall components. The complex biochemical mechanisms that build a microbial cell are not encoded by mammalian genomes, leading to cell wall components being considered PAMPs with no self-equivalent. Yet, even for PRRs that detect cell wall components, self-ligands have been reported. For example, the PRRs CD14, Toll-like Receptor 4 (TLR4), MD-2 and caspase-11 reportedly bind a collection of oxidized phosphatidylcholines (oxPAPC) that are produced by injured or dying mammalian cells (Chu et al. 2018; Erridge et al. 2008; Zanoni et al. 2017; 2016). Discussing the differences between self and nonself detection mechanisms by cell wall sensing PRRs is mainly a philosophical exercise, as we lack biochemical insight into how these receptor-ligand interactions occur.

In this study, we used the LPS and oxPAPC receptor caspase-11 as a model to define the mechanisms of self vs nonself detection. Using a variety of biochemical, bioinformatic, and cell-based assays, we discovered that despite their diversity of source, self and nonself lipids use a common mechanism to interact with caspase-11. This mechanism was defined at the level of the lipid and the ligand binding domain. The symmetry of mechanisms of interaction between self and nonself lipids extended to numerous caspase-11 homologs in distinct species of animals, knowledge of which enabled the derivation of an LPS receptor *de novo*. These findings provide important insight into the mechanisms of PAMP detection by caspase-11, and highlight self-ligand recognition as a (perhaps) unavoidable aspect of PRR activities.

## Results

### Distinct biochemical activities mediate caspase-11 binding and aggregation

Caspase-11 aggregation has served as an indicator of its interaction with LPS (Shi et al. 2014). This assay, however, has limitations, as LPS from *Rhodobacter sphaeroides* (LPS-RS) binds with high affinity to caspase-11 but does not induce aggregation (Shi et al. 2014). We verified these findings, as LPS-RS failed to induce caspase-11 aggregation on native PAGE (Figure 1B). To detect non-aggregating caspase-11 binding events, we utilized a competitive assay to inhibit caspase-11 aggregation in the presence of LPS from *E. coli* (LPS-EB, serotype O111:B4) and other competitive ligands. Co-incubation of caspase-11 with LPS-EB in the presence of increasing concentrations of LPS-RS hindered caspase-11 aggregation in this assay (Figure 1B). As expected (Shi et al. 2014), caspase-11 CARD alone demonstrates the ability to aggregate upon binding LPS-EB (Figure 1C).

**Figure 1:**
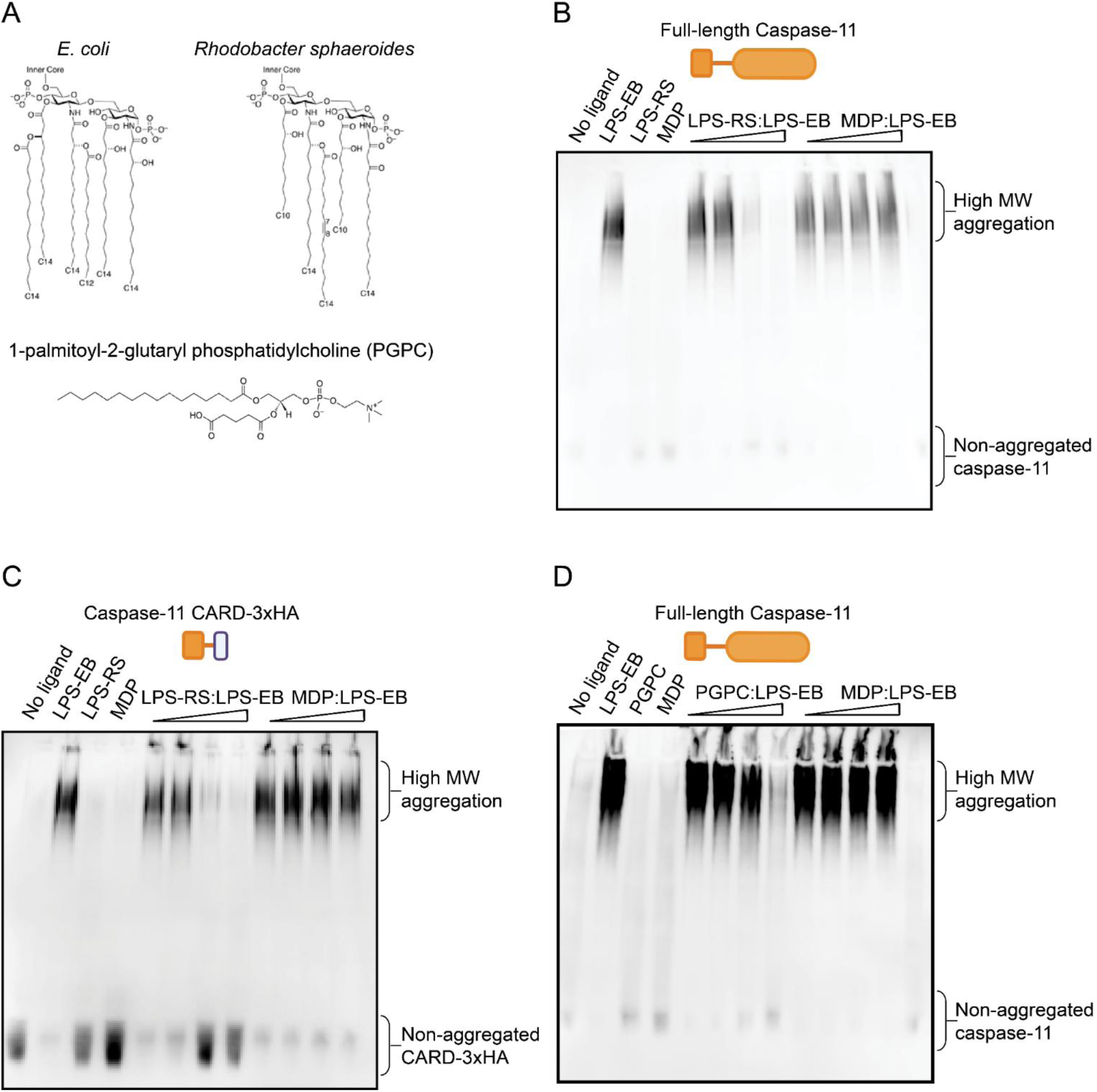
Caspase-11 CARD binds to monoacylated PGPC without aggregation. A) Chemical structures of lipopolysaccharides from *E. coli* (LPS-EB), *Rhodobacter sphaeroides* (LPS-RS), and 1-palmitoyl-2-glutaryl-sn-glycero-3-phosphocholine (PGPC). (B) 293T cell lysates expressing exogenous full-length caspase-11 were incubated at room temperature for 30 minutes with LPS-EB, LPS-RS, muramyl dipeptide (MDP), or combinations thereof. Protein complexes were resolved by blue native gel electrophoresis and analyzed via immunoblotting to detect caspase-11. (C) 293T cell lysates expressing HA-tagged caspase-11 CARD domainwere were incubated at room temperature for 30 minutes with LPS-EB, LPS-RS, muramyl dipeptide (MDP), or combinations thereof. Protein complexes were resolved by blue native gel electrophoresis and analyzed via immunoblotting to detect HA-tag. (D) 293T cell lysates expressing exogenous full-length caspase-11 werewere incubated at room temperature for 30 minutes with LPS-EB, PGPC, muramyl dipeptide (MDP), or combinations thereof. Protein complexes were resolved by blue native gel electrophoresis and analyzed via immunoblotting to detect caspase-11.

Like LPS, oxPAPC is a heterogenous mixture of lipids, which complicates biochemical interaction studies with caspase-11. Within the oxPAPC mixture of self-lipids is a chemically defined and soluble entity known as PGPC (1-palmitoyl-2-glutaryl phosphatidylcholine) (Zanoni et al. 2017). To determine if PGPC forms direct contacts with caspase-11, similar biochemical analyses were performed. We found that PGPC did not induce caspase-11 aggregation, as assessed by native PAGE, but PGPC inhibited LPS-induced caspase-11 and caspase-11 CARD aggregation (Figure 1D). PGPC may therefore bind caspase-11.

### An aggregation-independent assay to monitor lipid-caspase-11 interactions

We reasoned that caspase-lipid interactions would alter molecular conformations in a manner that reveals or obstructs protease cleavage sites. As such, limited trypsin digestion was used as a proxy for evaluating the interaction between caspase-11 and its ligands. Changes in molecular conformation may result in ligand-dependent tryptic digestion patterns that would be identified via immunoblot analysis (Figure 2A). This assay would therefore use differential trypic cleavage, rather than protein aggregation, as a readout of protein-lipid interactions.

**Figure 2:**
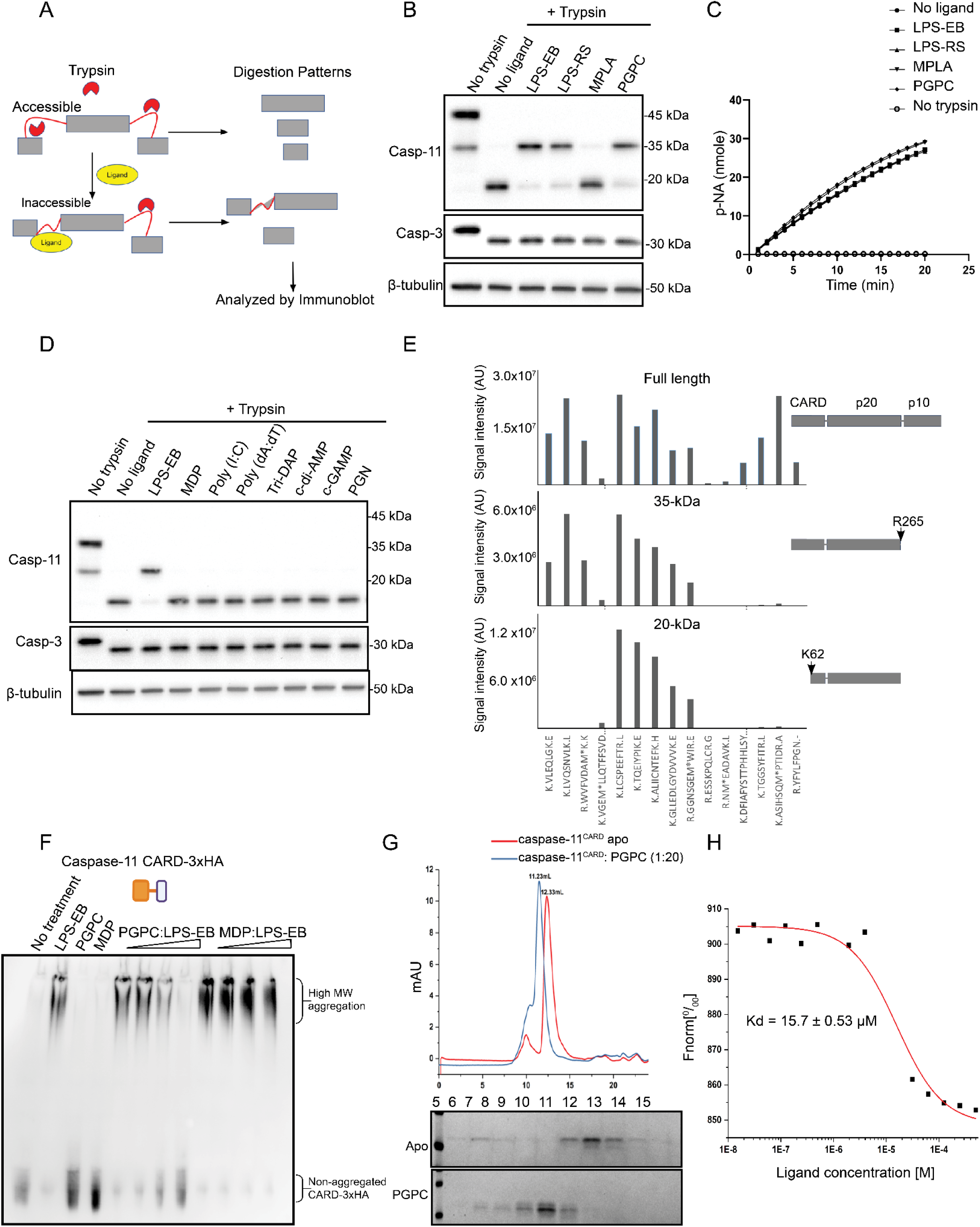
PGPC binds to the CARD of caspase-11. (A) Schematic representation of the principle of limited trypsin digestion. (B) 293T cell lysates containing caspase-11 were treated with various ligands, including LPS from *E. coli* (LPS-EB), *Rhodobacter sphaeroides* (LPS-RS), monophosphoryl lipid A (MPLA), and PGPC. Lysates were then subjected to limited trypsin digestion followed by immunoblot analysis to detect proteolytic cleavage of caspase-11. (C) Trypsin enzymatic activity in the presence of the indicated ligands was measured using a colorimetric assay, quantifying the impact of different ligands on trypsin enzymatic activity. (D) Caspase-11-containing cell lysates were treated with various bacterial ligands, including muramyl dipeptide (MDP), Poly(I), Poly(dA), Tri-DAP (L-Ala-γ-D-Glu-mDAP), c-di-AMP (cyclic di-adenosine monophosphate), cGAMP (cyclic guanosine monophosphate–adenosine monophosphate), and peptidoglycan (PGN). Limited trypsin digestion followed by immunoblotting was performed to assess ligand-induced proteolytic changes in caspase-11. (E) Full-length recombinant caspase-11 was incubated with PGPC, subjected to limited trypsin digestion, and analyzed by SDS-PAGE followed by Coomassie staining. Gel bands corresponding to full-length caspase-11 (and cleavage products of 35 kDa and 20 kDa) were further analyzed by mass spectrometry to identify cleavage sites. (F) 293T cell lysates expressing exogenous HA-tagged caspase-11 CARD with PGPC, LPS-EB, or MDP, alone or in combination, followed by blue native gel electrophoresis and immunoblotting for HA tag. (G) Recombinant caspase-11 CARD (amino acids 1–80) was incubated with PGPC with the molar ratio 1:20 and subjected to gel filtration chromatography. SDS-PAGE and Coomassie staining were used to analyze eluted fractions. (H) Microscale thermophoresis (MST) was performed to determine the binding affinity (Kd) between caspase-11 CARD (amino acids 1–80) and PGPC.

When exposed to trypsin, full-length (45 kDa) caspase-11 present within cell lysates underwent degradation within 15 minutes, yielding a 20 kDa fragment that was evident by immunoblot analysis (Figure 2B). In the presence of LPS-EB, caspase-11 exhibited a distinct state of trypsin resistance, resulting in the emergence of a higher molecular weight band (35 kDa) (Figure 2B). Similarly, LPS-RS and PGPC, which bind but do not aggregate caspase-11, exhibited similar banding patterns as LPS-EB during limited trypsin digestion (Figure 2B).

In contrast to the lipid-induced protection of caspase-11 to protease cleavage, the tryptic digestion patterns of caspase-3 and β-tubulin were unaffected by LPS-EB, LPS-RS, or PGPC (Figure 2B). Additionally, no alteration of intrinsic trypsin enzymatic activity (as assessed by spectrofluorimetry) was observed in the presence of lipids (Figure 2C). To assess the specificity of the trypsin resistance assay to report on caspase-ligand interactions, we examined caspase-11 cleavage in the presence of microbial products that are reported to not bind caspase-11 (Shi et al. 2014). In the limited trypsin digestion assay, we investigated the effect of various PAMPs including muramyl dipeptide (MDP), Poly(I:C), Poly(dA:dT), L-Ala-γ-D-Glu-mDAP (Tri-DAP), Bis-(3’-5’)-cyclic dimeric adenosine monophosphate (c-di-AMP), cyclic guanosine monophosphate–adenosine monophosphate (cGAMP), and peptidoglycan (PGN). None of these microbial products influenced the tryptic digestion of caspase-11 (Figure 2D).

To identify the regions of caspase-11 protected from tryptic digestion upon lipid binding, we conducted similar studies using recombinant caspase-11, followed by mass spectrometry. This analysis revealed that the 20 kDa band lacked peptides preceding K62, whereas the 35 kDa band retained the same N-terminal composition as the undigested full-length protein (Figure 2E). Notably, K62 resides within the CARD of caspase-11. Both the 35 kDa band (present only upon ligand binding) and the 20 kDa band (present in the absence of ligand) lacked peptides C-terminal to R265 (Figure 2E). These data suggested that binding of LPS or PGPC induces a common conformational change in the CARD that transforms it from a trypsin-sensitive to a trypsin-resistant state. Consistent with this idea, we found that PGPC prevented LPS-induced aggregation of the CARD of caspase-11 (Figure 2F). Additionally, liquid chromatography revealed that the caspase-11 CARD eluted in an earlier fraction in the presence of PGPC compared to caspase-11 CARD alone (Figure 2G). This shift in elution profile suggests a stable complex is formed between PGPC and the caspase-11 CARD. Microscale thermophoresis was employed to quantify the binding affinity between PGPC and the CARD, resulting in a determined Kd value of approximately 15 μM (Figure 2H). As the binding affinity between the caspase-11 CARD and PGPC is lower than the critical micelle concentration of PGPC (∼50μM) (Pande, Kar, and Tripathy 2010), the CARD of caspase-11 may have the capacity to interact with PGPC monomers.

### Caspase-11 binds lipids with diverse headgroup chemistries

Studies of LPS-CARD interactions have been complicated by the varied acylation and phosphorylation states of this PAMP. In contrast, PGPC offered a precisely defined chemical structure to explore the principles of caspase-11 interactions. Each PGPC molecule is comprised of a phosphatidylcholine headgroup, a 16-carbon acyl chain at the first carbon, and a glutaryl group at the second carbon (depicted in Figure 3A). Each of these structural features was examined for its impact on caspase-11 interactions. We first examined the glutaryl group in the interaction with caspase-11, through analysis of 16:0 LysoPC (1-palmitoyl-2-hydroxy-sn-glycero-3-phosphocholine). This lipid shares a comparable structure to PGPC, but lacks the glutaryl group at the second carbon (Figure 3A). Through the limited trypsin digestion assay, we observed that, similar to LPS and PGPC, 16:0 LysoPC protected caspase-11 from digestion, resulting in the 35 kDa protected band on immunoblot (Figure 3A). The glutaryl group is therefore not necessary to bind caspase-11, indicating that monoacylated lipids can interact with this protein.

**Figure 3:**
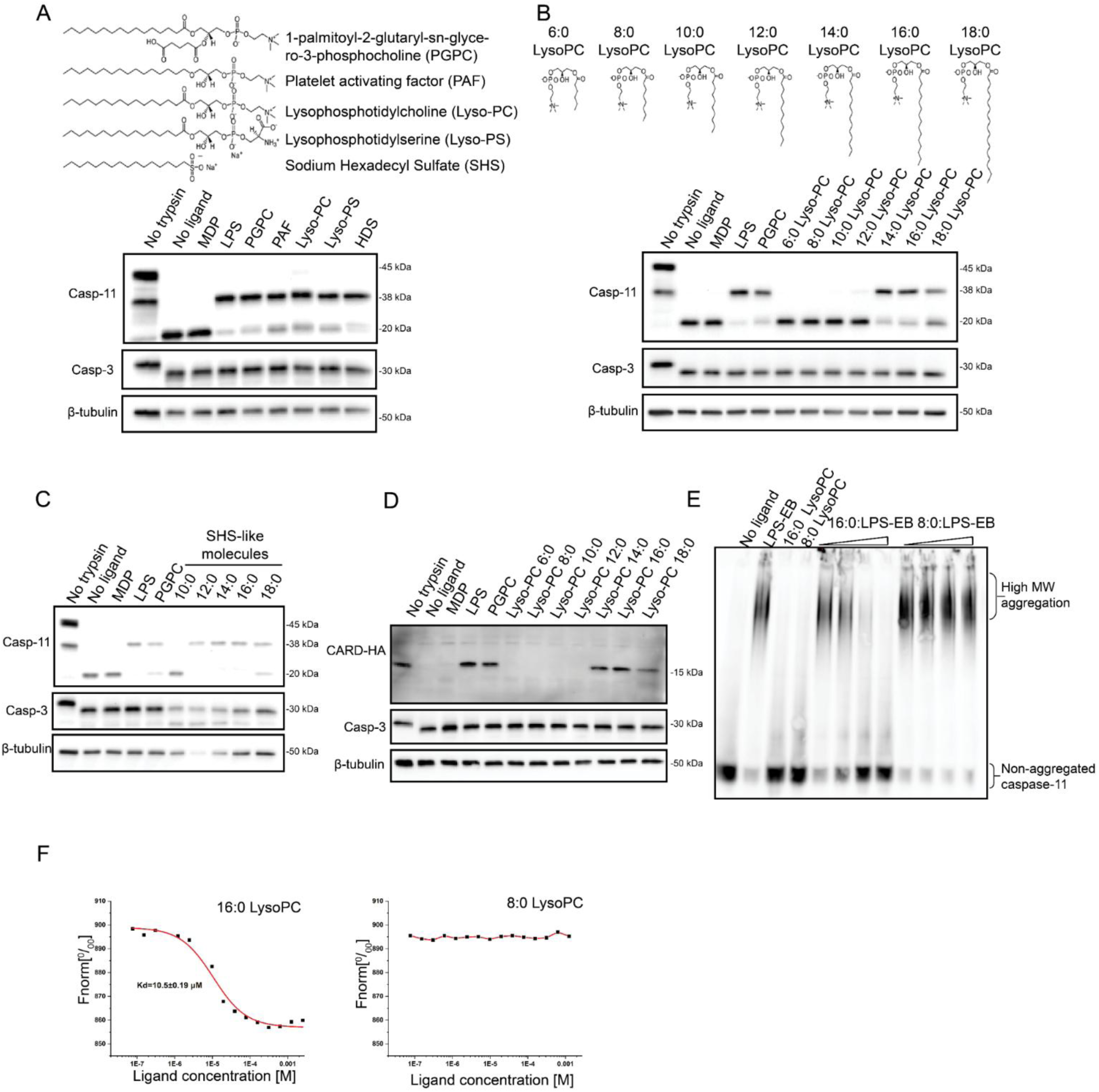
The binding capacity of lipids to caspase-11 CARD is determined by acyl chain length. (A) 293T cell lysates expressing caspase-11 were treated with various lipids featuring different headgroups, followed by limited trypsin digestion and immunoblotting to assess proteolytic sensitivity. (B) Caspase-11-containing cell lysates were treated with lysophosphatidylcholine (LysoPC) with varying acyl chain lengths, subjected to limited trypsin digestion, and analyzed by immunoblotting to detect proteolytic cleavage. (C) Cell lysates containing caspase-11 were treated with different sulfate surfactants of varying hydrocarbon chain lengths, followed by limited trypsin digestion and immunoblot analysis. (D) Lysates containing the caspase-11 CARD domain were treated with lipids of different headgroups, followed by limited trypsin digestion and immunoblot analysis. (E) Full-length caspase-11 cell lysates were treated with 16:0 or 8:0 LysoPCs, alone or in combination with LPS-EB, and analyzed by blue native gel electrophoresis followed by immunoblotting to detect caspase-11 aggregation. (F) The binding affinity (Kd) between caspase-11 CARD and 16:0 or 8:0 LysoPCs was determined using microscale thermophoresis (MST), highlighting differences in interaction strength between the lipid variants.

To determine the impact of the lipid headgroup on interactions with caspase-11, we performed experiments utilizing a series of 16-carbon monoacylated lipids. These lipids mainly encompassed a diverse range of negatively charged headgroups, spanning from those typically present in membrane phospholipids to the synthetic detergent sodium hexadecyl sulfate (Figure 3A). We found that most of the tested lipids displayed a protective effect on caspase-11 during limited trypsin digestion (Figure 3A), This finding suggests that caspase-11 can interact with monoacylated lipids containing diverse negatively charged headgroups.

### Acyl chain length is the primary determinant of caspase-11 interactions

Given the dispensability of the glutaryl group and the broad tolerance towards various headgroup chemistries, we explored the role of the acyl chain in facilitating caspase-11 interactions. Monoacylated lipids with one or two double bonds exhibited a protective effect on caspase-11 during limited trypsin digestion, suggesting that the degree of saturation in the acyl chain is not a determinant of caspase-11 binding (Supplementary Figure 1). We therefore explored the impact of acyl chain length on caspase-11 interactions. Limited trypsin digestion assays were performed in the presence of a spectrum of Lysophosphatidylcholines (LysoPCs) that contain acyl chains ranging from 6 carbons (6:0 LysoPC) to 18 carbons (18:0 LysoPC) (Figure 3B). These assays demonstrated that only LysoPCs possessing acyl chains of 14 carbons or longer protected caspase-11 under limited trypsin digestion (Figure 3B). Similar results were obtained with lipids having sulfate headgroups, where only lipids comprising a minimum of 12 continuous carbons provided protection against trypsin digestion (Figure 3C). The protective effect of these lipids was also observed in assays using the CARD of caspase-11, akin to our findings with LPS and PGPC. (Figure 3D).

To corroborate these findings, we examined lipid performance in the native PAGE caspase-11 aggregation assays described in Figure 1. We found that 16:0 LysoPC inhibited the aggregation caspase-11 induced by LPS-EB, whereas 8:0 LysoPC did not (Figure 3E). The findings suggested acyl chain length discrimination by caspase-11. Consistent with this idea, microscale thermophoresis demonstrated that 16:0 LysoPC displayed a binding affinity (Kd) comparable to PGPC (∼10 μM). In contrast, 8:0 LysoPC exhibited no detectable interaction with the caspase-11 CARD in this assay (Figure 3F). These findings indicate that the acyl chain length is a critical determinant governing the capacity of lipids to bind to caspase-11. In particular, lipids containing acyl chains 14 or more carbons in length are the preferred ligand for caspase-11.

### Amino acid determinants of caspase-lipid interactions

To determine if acyl chain binding is a conserved feature across mammals, we cloned the CARDs of caspase-11 orthologues (annotated as caspase-4) from various mammalian species and subjected them to limited trypsin digestion. Under the same conditions used for mouse caspase-11, PGPC was found to protect the 16 out of 22 tested CARDs from tryptic digestion. Notably, this protection was observed even in CARDs that showed no protection when exposed to LPS under the same conditions used to detect LPS and PGPC binding to mouse caspase-11 (Figure 4A). Acyl chain binding may therefore be a more common feature among mammalian caspase-4 proteins than LPS binding activities. Aligning the sequences that exhibited PGPC protection and comparing them to those that were not protected revealed two conserved phenylalanine residues, Phe45 and Phe76, that were abundantly present in the PGPC-protected group (Figure 4B). Neither of these amino acids have been identified as determinants of caspase-11 function. Although not investigated further herein, we noted that human caspase-5 and the caspases from sheep and lemur do not contain both Phe45 and Phe76, but are still protected by PGPC. This finding suggests that other residues may contribute to acyl chain binding.

**Figure 4:**
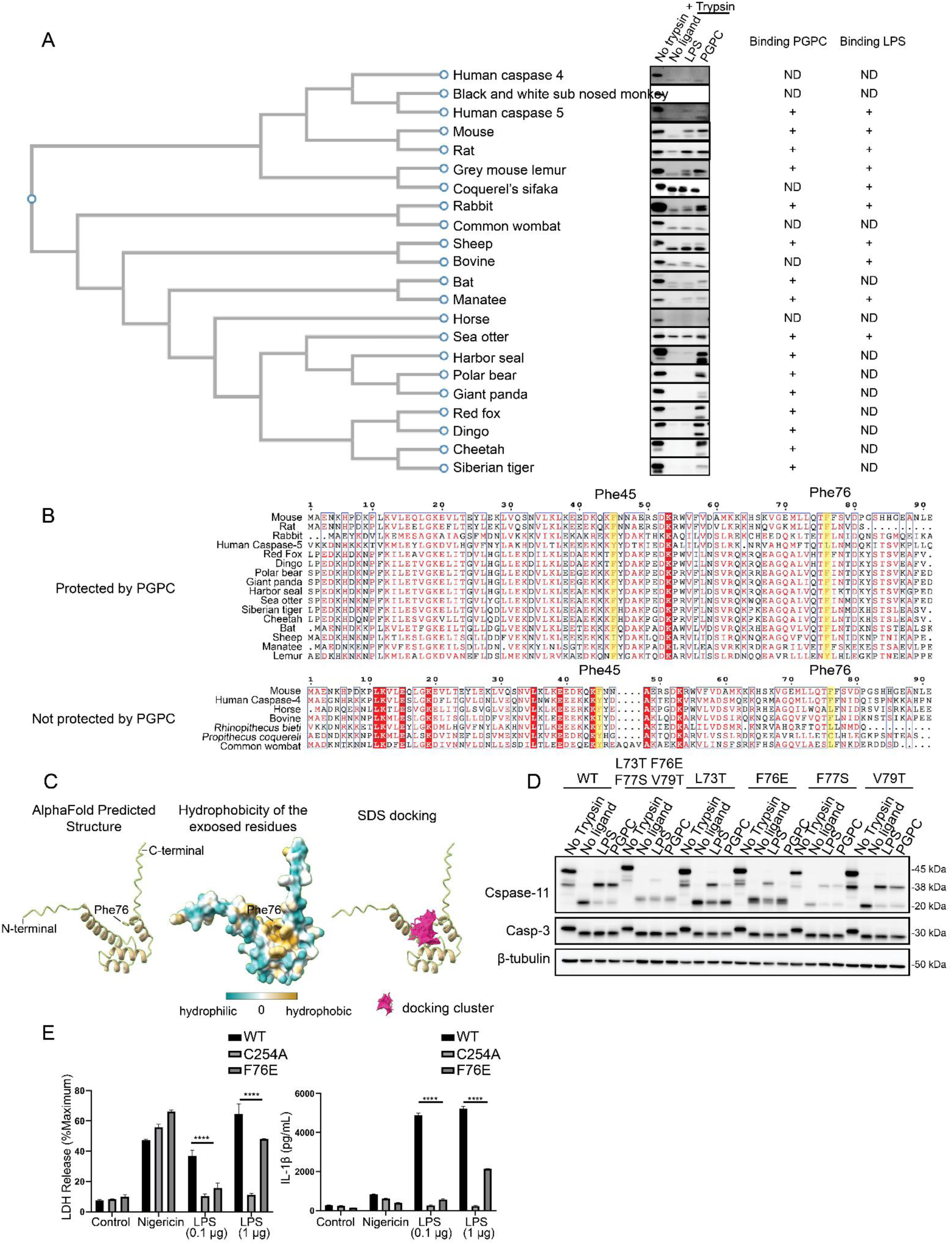
Structural and functional analysis of caspase-11 CARD homologs and key residue mutants. (A) Various mammalian caspase-11 CARD orthologs were exogenously expressed in 293T cells. Cell lysates were incubated with PGPC (100 µg/mL) and LPS (10 µg/mL), followed by limited trypsin digestion. The outcomes were categorized based on a phylogenetic tree constructed using ClustalW. (B) Amino acid sequence alignment of caspase-11 CARD domains from different orthologs, grouped by protection or lack of protection from PGPC. Conserved residues, Phe45 and Phe76, are highlighted in the protected group. (C) AlphaFold-predicted structure of caspase-11 CARD, showing a hypothetical hydrophobic pocket, predicted to accommodate the hydrocarbon chain of sodium dodecyl sulfate (SDS)by SwissDock. The conserved Phe76 is predicted to be at the bottom of the hypothetical hydrophobic pocket. (D) Site-directed mutagenesis of hydrophobic residues in the 70–80 region of caspase-11 CARD, including the F76E mutation were expressed in 293T cells. Cell lysates were incubated with PGPC (100 µg/mL) and LPS (10 µg/mL), followed by limited trypsin digestion and immuneblot to detect caspase-11. (E) Lactate dehydrogenase (LDH) and IL-1β release in caspase-11 knockout immortalized macrophages reconstituted with caspase-11 variants including the F76E mutation. Cells were electroporated with LPS to assess the functional impact of the CARD variants on cytokine release and cell death. Data are representative of three independent experiments, and results with error bars are presented as means ± SEM. Statistical significance was determined by an ordinary two-way ANOVA compared to AcCARD. ns = not significant, *p<0.05, **p<0.01, ***p<0.001, ****p<0.0001

*In silico* analysis was employed using AlphaFold (Mirdita et al. 2022) to model the CARD-lipid interaction. This analysis suggested that the CARD has a configuration featuring four amphipathic alpha helices, deviating from the conventional six helices found in canonical members of the death domain superfamily (Figure 4C) (Park et al. 2007), and is consistent with the reported crystal structure of MBP-tagged caspase-11 CARD tetramer (M. Liu et al. 2020). The CARD may also contain a hydrophobic cleft, partly exposed to the solvent (Figure 4C). To probe the potential of this cleft to accommodate acyl chains, we conducted molecular docking utilizing Swissdock (Grosdidier, Zoete, and Michielin 2011b; 2011a). This analysis focused on the binding interaction with sodium dodecyl sulfate (SDS), the smallest sulfate-containing lipid that protect caspase-11 under limited trypsin digestion (Figure 4C). The docking outcomes revealed a predominant location of SDS within the hydrophobic cleft of the CARD (Figure 4C). Phe76 is positioned within the hydrophobic pocket predicted by AlphaFold (Figure 4C), suggesting a role in lipid binding. Mutating Phe76 to glutamic acid abolished the trypsin protection induced by PGPC and diminished the protective effect of LPS on tryptic digestion (Figure 4D).

To assess the functional impact of these *in vitro* findings on caspase-11 activity within cells, we reconstituted caspase-11-deficient immortalized bone marrow-derived macrophages (iBMDMs) with wild-type (WT), catalytically inactive (C254A), and acyl chain binding-deficient (F76E) caspase-11 variants. These cells were then subjected to LPS delivery via electroporation. As expected (Kayagaki et al. 2011; Shi et al. 2014)), cells expressing WT caspase-11 responded to electroporated LPS by releasing the cytoplasmic proteins IL-1β and lactate dehydrogenase (LDH) into the extracellular space. Also as expected (Kayagaki et al. 2011; Shi et al. 2014), cells expressing the C254A mutant were defective for these LPS responses (Figure 4E). Consistent with the results from limited trypsin digestion, cells reconstituted with the caspase-11 F76E variant were defective in LPS-induced LDH and IL-1β release at low LPS concentration, with partial responses observed at higher LPS doses. Importantly, all three cell lines displayed similar LDH release and IL-1β production upon exposure to LPS and nigericin, a stimulus that functions independently of caspase-11 (Kayagaki et al. 2011) (Figure 4E). Phe76, an amino acid we first identified based on self-ligand (PGPC) interaction studies, is therefore required to bind LPS *in vitro* and to mediate LPS-induced inflammatory activities within cells.

### Two functional regions within the CARD of caspase-11 mediate lipid interactions

While Phe76 is required for interactions between caspase-11 and LPS or PGPC, the outcome of these interactions is not identical. LPS-induces caspase-11 oligomerization whereas PGPC does not (Figure 1). Additional regions within the CARD may therefore mediate the transition from lipid binding to oligomerization. To explore this concept, we examined the N-terminal region of the CARD, upstream of Phe45 and Phe76. A series of N-terminal truncated murine caspase-11 mutants was generated and assessed for binding to LPS and PGPC using limited trypsin digestion and native PAGE. The first ten amino acids proved dispensable for both LPS and PGPC binding. Removal of amino acids 10-15 abolished caspase-11 aggregation (Figure 5A) and LPS binding (Figure 5B), while the trypsin protective effect induced by PGPC was retained (Figure 5B). Binding to PGPC was only compromised upon the removal of amino acids 20-25 (Figure 5B). This observation suggests a model in which two functionally distinct regions coexist within the caspase-11 CARD: 1) a C-terminal region responsible for acyl binding, which is independent of aggregation, and 2) an upstream N-terminal region that also contributes to lipid binding but uniquely facilitates protein aggregation.

**Figure 5:**
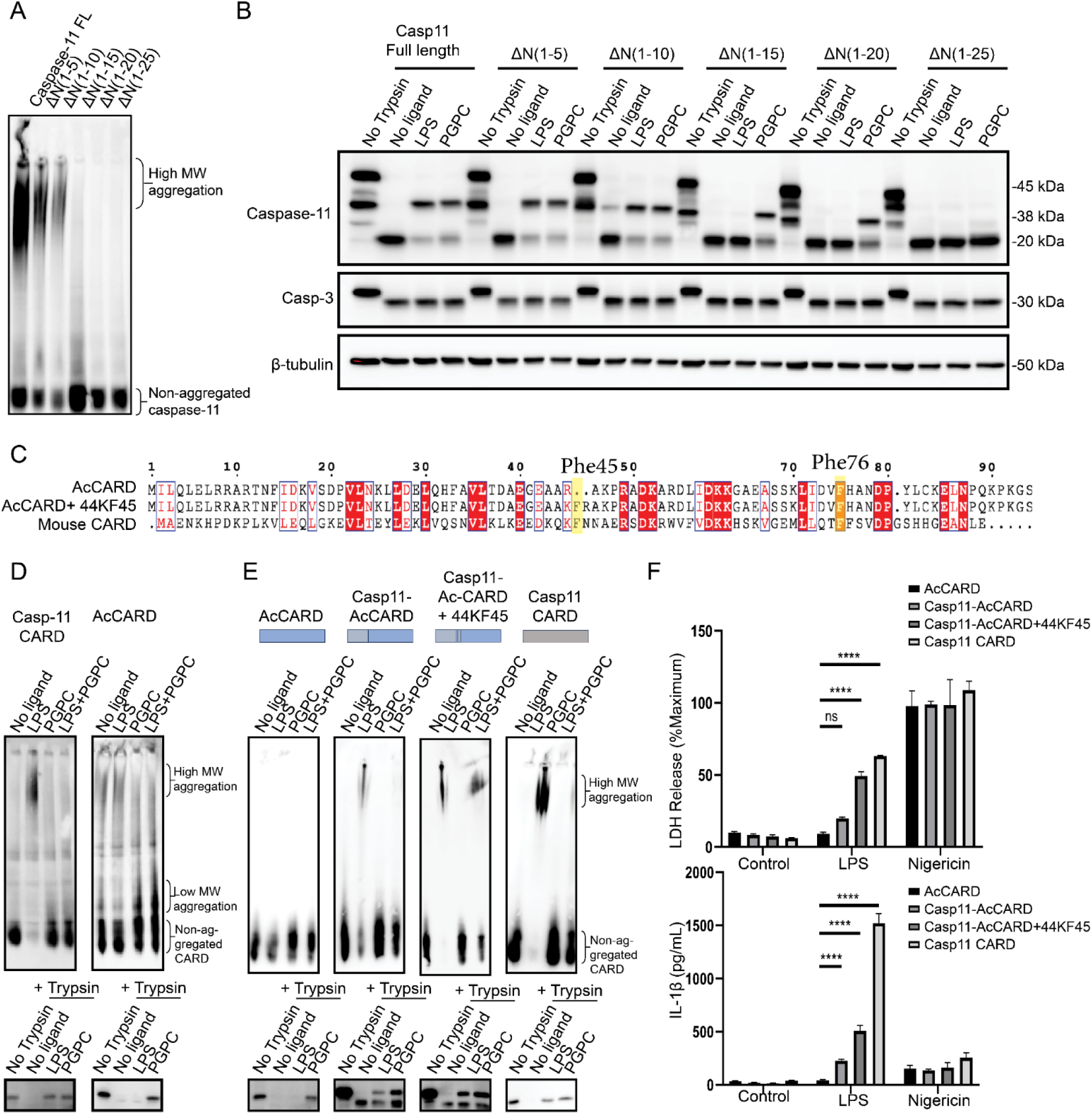
The N-terminal region of caspase-11 mediates caspase-11 aggregation by LPS. (A) 293T cell lysates expressing N-terminal truncated mutants of caspase-11 were incubated with LPS (10µg/mL) for 30 minutes at room temperature. The lysates were analyzed by blue native PAGE followed by immunoblotting to detect caspase-11. (B) 293T cell lysates expressing N-terminal truncated mutants of caspase-11 were incubated with LPS (10µg/mL) and PGPC (100µg/mL) for 30 minutes at room temperature and subjected to limited trypsin digestion, followed by immunoblotting to detect caspase-11. (C) Sequence alignment comparing the CARD domains of *Amphilophus citrinellus* CARD-containing protein (AcCARD), mouse caspase-11 CARD, and AcCARD with lysine and phenylalanine added at positions 44 and 45 (AcCARD+44KF45). (D) 293T cell lysates expressing HA-tagged AcCARD and caspase-11 CARD were incubated with LPS (10µg/mL) and PGPC (100µg/mL) for 30 minutes at room temperature and then subjected to limited trypsin digestion and blue native PAGE followed by immunoblot to detect HA. PGPC induced AcCARD aggregation is indicated by (E) 293T cell lysates expressing engineered HA-tagged AcCARD constructs, where the first 38 amino acids on the N-terminus were replaced by the corresponding residues of caspase-11 CARD (Casp11-AcCARD) and an additional variant with K44 and F45 added (Casp11-AcCARD+44KF45), were incubated with LPS (10µg/mL) and PGPC (100µg/mL) for 30 minutes at room temperature. These lysates were then analyzed by blue native PAGE and limited trypsin digestion, followed by immunoblotting to detect HA. (F) IL-1β and LDH release were measured in caspase-11 knockout immortalized macrophages reconstituted with chimeric proteins, where the caspase-11 catalytic domain was fused to different CARD variants. Cells were electroporated with LPS to assess the functional impact of the CARD variants on cytokine release and cell death. Data are representative of two independent experiments, and results with error bars are presented as means ± SEM. Statistical significance was determined by an ordinary one-way ANOVA compared to AcCARD. ns = not significant, *p<0.05, **p<0.01, ***p<0.001, ****p<0.0001

If this model is correct, then the two functionally distinct activities within the CARD may be dissociable, and therefore amenable to bioengineering-based manipulation. To test this idea, we sought to engineer CARD-like LPS binding activities within proteins that do not naturally have such functions. Such bioengineering events have been used to rewire the effector domains present in innate immune signal transduction pathway components (Devant, Cao, and Kagan 2021; Tan and Kagan 2019). However, it is unclear if a PRR can be engineered to change ligand specificity. To explore this idea, we first identified additional CARDs that lacks LPS binding activity. We utilized the Foldseek Search Server (Van Kempen et al. 2023), which identifies proteins that share similar tertiary structures and amino acid interactions as that predicted for the caspase-11 CARD by AlphaFold. This *in silico* analysis identified an uncharacterized CARD-containing protein (Uniprot A0A3Q0SFM2) in cichlid fish (*Amphilophus citrinellus*) (hereafter AcCARD) (Figure 5C). Biochemically, we found that AcCARD binds PGPC in both native PAGE and limited trypsin digestion assays. AcCARD showed no evidence of LPS binding (Figure 5D). PGPC induced some degree of higher molecular weight AcCARD complexes, by a process that was not inhibited by co-treatment with LPS (Figure 5E). AcCARD therefore represents a specific PGPC-binding protein, with no ability to bind LPS.

Based on these data, we determined if we could convert AcCARD into an LPS receptor. Sequence alignment between caspase-11 CARD and AcCARD revealed a relatively conserved sequence in the C-terminus. However, the N-terminal residues displayed opposite charges, with positively charged residues in caspase-11 aligning with negatively charged residues in AcCARD. To determine if the N-terminal region of the caspase-11 CARD could confer LPS-binding activity to AcCARD, we fused the first 38 amino acids of caspase-11 CARD to the PGPC-binding domain of AcCARD (amino acids 39-90). This hybrid protein (Casp11-AcCARD) acquired the ability to aggregate and demonstrated protection under limited trypsin digestion in the presence of LPS (Figure 5E). Furthermore, reconstitution of caspase-11 deficient immortalized macrophages with the hybrid Casp11-AcCARD fused to the catalytic domain of caspase-11 resulted in cell death and IL-1β release in response to LPS electroporation (Figure 5F). The efficacy of this *de novo* LPS receptor to induce pyroptosis was further enhanced by incorporating two additional amino acids at position 44 and 45, including the conserved Phe45 found in PGPC-protected caspase 4 CARDs (Casp11-AcCARD 44KF45) (Figure 5E, F).

Close inspection of caspase-11 amino acids 26 to 30 revealed a lysine at position 28, which was absent in the corresponding sequence of AcCARD. Introducing this single substitution into AcCARD (E29K) was sufficient to convert AcCARD into an LPS-binding domain, as demonstrated by both aggregation assays and limited trypsin digestion (Figure 6A). Furthermore, the LPS-induced aggregation and protective effects were enhanced when an additional mutation at position 20 (D20K), corresponding to the conserved K19 in caspase-11, was introduced (Figure 6A). Thus, substitution of two amino acids in AcCARD is sufficient to convert this protein from a PGPC receptor into a dual PGPC/LPS receptor, akin to caspase-11. The importance of these amino acids (K19 and K28) in LPS responses by murine caspase-11 was confirmed in reconstituted caspase-11 deficient macrophages. Cells expressing K19, K28 double mutations were unresponsive to electroporated LPS but retained the ability to respond to LPS + nigericin (Figure 6B). However, while AcCARD D20K/E29K displayed LPS-binding activity *in vitro*, cells reconstituted with AcCARD D20K/E29K fused to the caspase-11 catalytic domain failed to respond to electroporated LPS (Figure 6C). This failure may be attributed to spontaneous auto-cleavage of the mutant proteins within macrophages, resulting in nonfunctional caspase-11 fragments, which warrants further investigation (Figure 6D). Nevertheless, the ability to convert a PGPC-binding protein into an LPS receptor *in vitro* and within cells using chimeric approaches supports the existence of two loosely coupled regions within CARDs. The C-terminal region mediates lipid interactions through hydrophobic interactions, while the upstream N-terminal region uses positively charged residues to link LPS binding to protein aggregation.

**Figure 6:**
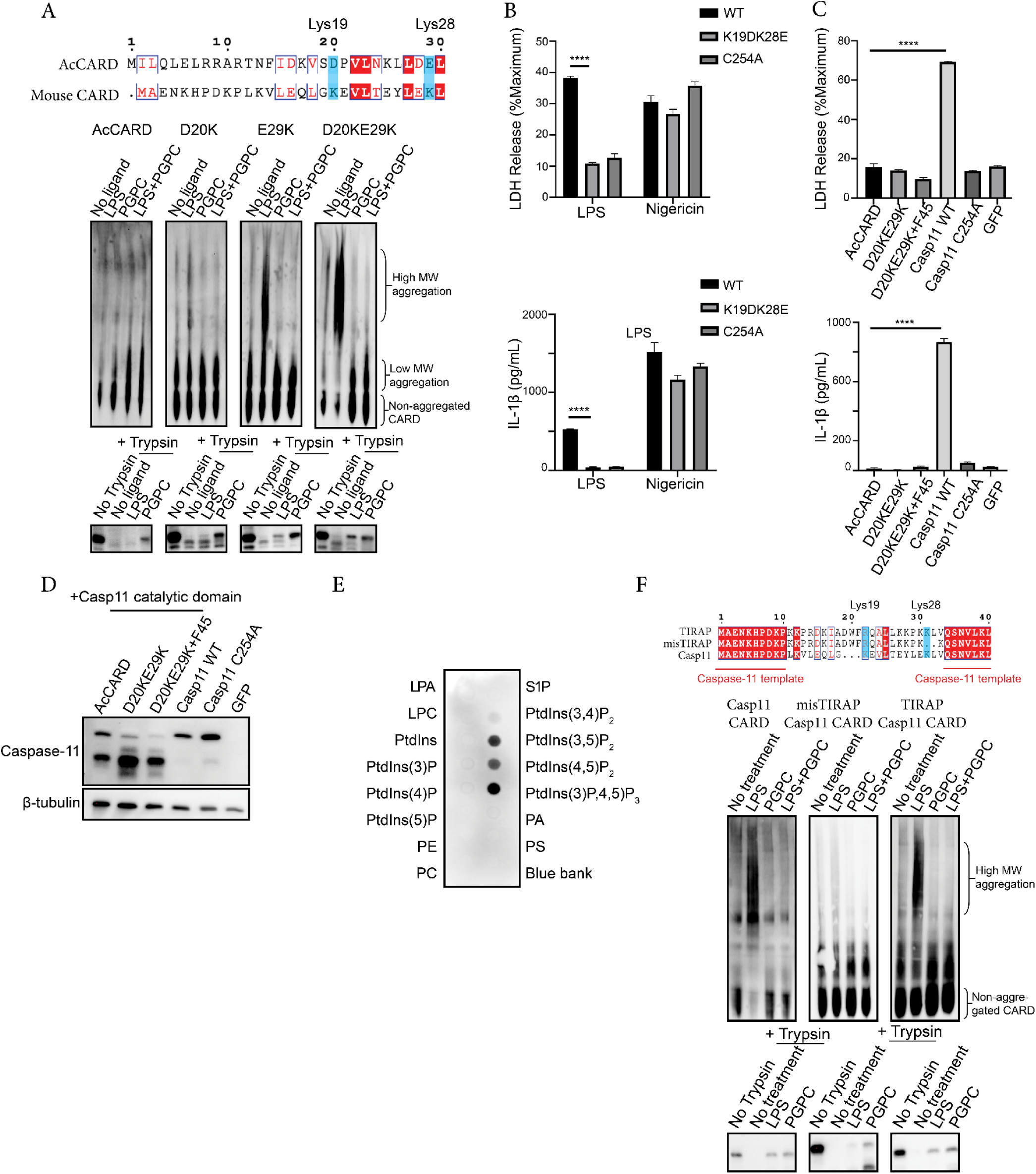
Caspase-11 binds to phosphoinositides. (A) 293T cell lysates expressing HA-tagged AcCARD with mutations converting negatively charged residues to the corresponding positively charged residues in caspase-11 CARD were incubated with LPS (10 µg/mL) and PGPC (100 µg/mL) for 30 minutes at room temperature. Lysates were subjected to limited trypsin digestion and blue native PAGE, followed by immunoblotting to detect HA. (B) IL-1β and LDH release were measured in caspase-11 knockout immortalized macrophages reconstituted with the K19DK28E mutant and the catalytically inactive mutant C254A. Cells were electroporated with LPS (1 µg per 10⁶ cells) to assess the functional impact of these CARD variants on cytokine release and cell death. Data are representative of three independent experiments, and results with error bars are presented as means ± SEM. Statistical significance was determined using one-way ANOVA compared to AcCARD. *ns = not significant, *p < 0.05, **p < 0.01, ***p < 0.001, ***p < 0.0001. (C) IL-1β and LDH release in caspase-11 knockout immortalized macrophages reconstituted with the caspase-11 catalytic domain fused to AcCARD variants (D20KE29K, D20KE29K+44KF45), followed by LPS electroporation. Data are representative of three independent experiments, and results with error bars are presented as means ± SEM. Statistical significance was determined by an ordinary one-way ANOVA compared to AcCARD. ns = not significant, *p<0.05, **p<0.01, ***p<0.001, ****p<0.0001 (D) Immunoblot analysis of caspase-11 fused to different AcCARD variants at the basal state, showing protein expression levels and complex formation. (E) Recombinant caspase-11 was incubated with a PIP-strip containing various phosphoinositides, followed by immunoblot detection to assess phosphoinositide binding by caspase-11. (F) 293T cell lysates expressing engineered CARDs containing caspase-11 N-terminal residues (30-90) fused to the TIRAP phosphoinositide-binding motif (TIRAP Casp11 CARD) and a misaligned control (misTIRAP Casp11 CARD) were incubated with LPS (10 µg/mL) and PGPC (100 µg/mL). Samples were analyzed by blue native PAGE and limited trypsin digestion, followed by immunoblotting to detect caspase-11 interactions.

### Caspase-11 binds to phosphoinositides

Given the role of N-terminal positively charged residues in mediating LPS-binding and aggregation, we reasoned that the corresponding molecular cue in caspase-11 ligands possesses a negative charge. LPS contains a negatively charged group, in the form of the bis-phosphorylated glucosamine component of lipid A. We reasoned that if lipid headgroup charge was a determinant of caspase-11 interactions, then other phosphorylated lipids should display evidence of caspase-11 interactions. To explore this concept, we examined the interactions between caspase-11 and a variety of eukaryotic lipid species. We employed a PIP-strip assay, a method where lipids are absorbed onto a hydrophobic membrane and subsequently incubated with recombinant protein (Shirey, Scott, and Stahelin 2017). We found that recombinant caspase-11 bound to diverse phosphorylated lipids, including PI(3,4,5)P3 and several phosphatidylinositol bisphosphates (Figure 6E), which are common constituents of eukaryotic cellular membranes (Posor, Jang, and Haucke 2022). No detectable interaction was observed with lipids or sphingolipids that lack phosphorylated headgroups (Supplementary Figure 2). These findings expand the repertoire of self lipids caspase-11 binds beyond PGPC, and suggest preferentially interactions with phosphoinositides with multiple phosphate groups (akin to what is present in bis-phosphorylated LPS). We note that caspase-11 did not bind to Lyso-PC on the strip (Supplementary Figure 2), likely because the acyl chain of these lipids was absorbed into the hydrophobic membrane, leaving only the headgroup available for interaction with caspase-11. This finding supports the general belief that PIP strips report on lipid headgroup interactions, whereas the limited trypsin digestion assay reports on acyl chain interactions.

Proteins interact with phosphoinositide headgroups through electrostatic interactions, whereby positively charged amino acids in a lipid binding domain interact with the negatively charged phosphate groups (Posor, Jang, and Haucke 2022). We reasoned that if a similar principle applies to CARD-lipid interactions, then swapping the N-terminal region of caspase-11 with a heterologous phosphoinositide-binding domain should maintain LPS interactions. We therefore engineered a fusion protein between the phosphoinositide-binding domain of Toll-like receptor adaptor protein TIRAP (Kagan and Medzhitov 2006) and the C-terminal acyl chain-binding region of caspase-11 CARD. Two chimeric proteins were engineered. One was generated by aligning the positively charged residues in the phosphoinositide-binding domain of TIRAP with the conserved N-terminal positively charged residues in caspase-11 CARD (TIRAP-CARD). A second chimeric protein did not utilize this alignment to position positively charged residues, referred to herein as misTIRAP-CARD (Figure 6F). We found that the TIRAP-CARD fusion protein exhibited signatures of LPS-binding, including aggregation and protection under limited trypsin digestion in the presence of LPS (Figure 6F). In contrast, misTIRAP-CARD, which lacked proper alignment of positively charged residues, did not exhibit evidence of LPS interactions (Figure 6F). Binding to negative charged headgroups, present on either self (PIPs) or nonself (LPS) lipids, is therefore sufficient to link acyl chain binding to aggregation of the CARD of caspase-11.

## DISCUSSION

Previous research has described the roles of the phosphate group and the hydrophobic component within lipid A in caspase interactions (Chen et al. 2023; Chu et al. 2018; Shi et al. 2014). In this study, our analysis expanded these prior conclusions beyond LPS to include self lipids with similar properties, thereby revealing the biochemical essence of a caspase-11 ligand. We propose that the caspase-11 ligand is not LPS specifically, but rather any lipid that contains two molecular cues: acyl chains with 14 or more carbons and a negatively charged head group. The N-terminal region of the caspase-11 CARD interacts with the charged lipid headgroup, whereas the C-terminal region mediates acyl chain interactions. Lipids that contain both of these features (*e.g.* LPS) enable the conversion of caspase-11 from an inactive monomer to an active oligomer that drives pyroptosis and inflammation. Lipids that contain one of these features, such as PGPC monomers, can bind but not aggregate caspase-11 efficiently. While structural analysis will be required to provide unimpeachable evidence to support this model, the strongest functional evidence to support the idea of a bipartite LPS-binding CARD comes from the chimeric proteins we generated. The most notable of these converted a PGPC-binding protein (AcCARD) into an LPS receptor (*in vitro* and in cells). Further support for this model came from the LPS-binding chimera where the N-terminal caspase-11 region was swapped with the PIP binding domain of TIRAP.

Of the amino acids identified that mediate lipid interactions, K19 was previously shown to be essential for caspase-11 to bind LPS (Shi et al. 2014). While K19 is conserved across all caspase-4 homologs, we identified residue, K28, which is uniquely present in rodents (Figure 4B). This amino acid appears to be a key determinant of aggregation, as K28 is required for LPS-induced aggregation of caspase-11, and introducing K28 into AcCARD was sufficient to confer LPS aggregation. We propose that K28 accounts for the ease at which LPS interactions with caspase-11 can be observed, as compared to other caspases whose LPS binding activity is less easy to detect (*e.g.* human caspase-4).

It has not escaped our attention that a pivotal question arises from our finding that self lipids display the key features associated with LPS-caspase interactions. Is self-ligand detection a mistake or a beneficial product of evolution? We consider this question from two perspectives. The first perspective on the role of self-ligand detection is cell-based, and highlights self-ligand detection as a damage-sensing mechanism that may be beneficial for host defense. Hydrophobic moieties, such as acyl chains with 14 or more carbons, are typically absent in the cytosol. These moieties are usually embedded within the hydrophobic regions of membranes or sequestered in lipid droplets, making them inaccessible to cytosolic caspase-11. Therefore, long acyl chains in the cytosol may act as damage-associated molecular patterns (DAMPs) detectable by caspase-11, leading to activities that may promote repair and a return to homeostasis. Such activities for caspase-11, if they exist, are undefined.

In addition to freely exposed acyl chains in the cytosol, it is proposed that certain lipid compositions within cellular membranes may disrupt the typical lamellar structure, leading to the exposure of acyl chains that are normally concealed within the bilayer core (Scollo et al. 2024). The possibility that acyl chains can be exposed on cellular membranes suggests a context-dependent mechanism for the regulation of caspase-11 binding and activation. Given the requirement for proximity and auto-processing to achieve full activation (Ross et al. 2018), if these hypotheses hold true, this scenario illustrates a three-dimensional threat-associated molecular pattern—responding to alterations in both the composition and topology of cellular membranes, potentially triggered by pathogen invasion or cellular stress. Furthermore, it is possible that caspase-11 binds phosphoinositides in a physiological context, which would enable interactions with intracellular vesicles, positioning this PRR as a guardian against bacterial escape from phagosomes.

The second perspective on the role of self ligand detection relates to self detection as a guardrail to prevent dysfunctional PRR evolution. A key feature of PRRs is their ability to recognize conserved molecular patterns associated with microbial products (*i.e.* PAMPs). This function must remain stable to ensure their effectiveness in immune defense. However, foreign molecular patterns are not ideal for maintaining consistent selective pressure for three key reasons. First, the same molecular pattern can vary significantly across microbial species, strains, or physiological conditions even for a conserved component like lipid A of LPS (Dardelle et al. 2024; Maldonado, Sá-Correia, and Valvano 2016). Second, threats are rarer and less predictable than non-threats, with the most successful pathogens often evolving to evade PRRs (Kagan 2023). Lastly, there is significant redundancy in threat detection, with multiple PRRs having the ability to detect individual microbes. This redundancy could enable genetic drift, allowing PRRs to lose their capacity for detection of the original molecular pattern they evolved to detect.

We propose that self ligands for PRRs may serve as a stable benchmark to preserve the ability of PRRs to recognize molecular patterns effectively. In this regard, it is possible that self ligand detection by PRRs is not an activity that is counter-selected, but rather a fundamental feature that is needed to ensure the biochemical essence of the target ligand remains unchanged throughout evolution. We note that, if this proposed self first model of PRR function is correct, significant regulatory mechanisms would need to be in place to ensure homeostasis and the induction of inflammation when needed. Such regulatory mechanisms are already evident in the context of nucleic acid sensing PRRs, as several nucleases and other enzymes are needed to prevent detection of self DNA and RNA (Crow et al. 2015; Jimeno et al. 2021; Rodero et al. 2017; Stetson et al. 2008; Uggenti, Lepelley, and Crow 2019). Whether analogous mechanisms exist for non-nucleic acid sensing PRRs is presently unclear. The concepts described herein provide a mandate to expand the studies of self ligand detection to all members of the PRR superfamily.

## MATERIALS AND METHODS

### Gene cloning

Genes of interest following the Kozak sequence (GCCACC) were cloned into the pMSCV-IRES-GFP vector between the XhoI and NotI restriction sites. A triple HA-Tag was added to the C-terminus of all CARD structures, linked by a triple glycine-serine (GC) linker (GGGGSGGGGSGGGGSG).

### 293T transfection

293-T cells were seeded 12-16 hours prior to transfection in a 6-well plate at a density of 7.5 x 10^5^ cells in a volume of 2 ml per well. On the day of transfection, 2.5 µg of plasmid DNA was diluted in 100 µL of OptiMEM and mixed with 7.5 µL of Lipofectamine in 100 µL of OptiMEM. The Lipofectamine mixture was incubated at room temperature for 15-20 minutes and then combined with the plasmid DNA. The transfection mixture was added dropwise to the wells containing the cells and incubated at 37 °C for 16-20 hours.

### Cell lysate preparation

On the day following transfection, cell lysates were prepared by adding 200 µL of ice-cold lysis buffer (150 mM NaCl, 25 mM Tris (pH 7.5), and 5 mM MgCl₂, and 0.05% Triton X-100) to each well. Cells were scraped using a sterile cell scraper, and lysates were vortexed intermittently while incubating on ice for 20 minutes. Lysates were centrifuged at 16,000g for 5 minutes at 4 °C, and the supernatant was collected. Lysates (140 µL, ∼600 µg of total protein) were aliquoted into a 96-deep-well plate. To each well, 5 µL of LPS (1 mg/mL, ALX-581-012) or 50 µL of lipids (1 mg/mL) were added, and the total volume was brought to 500 µL with LTD buffer (150 mM NaCl, 25 mM Tris (pH 7.5), and 5 mM MgCl₂). The mixture was incubated at room temperature for 30 minutes.

### Limited trypsin digestion

Trypsin (Sigma Aldrich, T1426) was freshly prepared by weighing 10 mg of trypsin powder and resuspending it in DMEM to a concentration of 1 mg/mL. This was diluted to a 10X concentration of 50 µg/mL with LTD buffer. 50 µL of lysate were removed from each well as a time zero control. 50 µL of diluted trypsin (50 µg/mL) were added to each well, mixed thoroughly, and incubated. Samples were collected at specific time points, and 50 µL of each sample was added to an equal volume of 2X Laemmli buffer followed by heating at 95°C for 5 minutes to stop the reaction. For caspase-11 in 293-T cell lysates, a 15-minute time point was optimal. Samples were then subjected to SDS-PAGE and immunoblot analysis (20 µL per well).

### Blue native PAGE

10 µl of lysate were incubated with 1 µg of O111:B4 LPS (1 mg/mL, ALX-581-012) and 10 µg of lipids (1 mg/mL, ALX-581-012), with the volume adjusted to 20 µL using ultrapure water. The mixture was incubated at room temperature for 20 minutes. NativePAGE sample buffer (Thermo Fisher Scientific, BN2003) was then added to the reaction mixture. The samples were loaded onto NativePAGE™ 4-16% (Thermo Fisher Scientific, BN1004BOX) Bis-Tris Protein Gels (1.0 mm, 15-well) and run according to the protocol for immunoblot analysis, using dark blue buffer (Thermo Fisher Scientific, BN2002) for the first third of the gel and light blue buffer for the remainder. The gel was subsequently transferred to a PVDF membrane for immunoblot analysis.

### Competitive binding assay

10 µl of lysate were incubated with 1 µg of O111:B4 LPS (1 mg/mL, Enzo ALX-581-012) and increasing concentrations of LPS-RS (0.5, 1, 5, 10 w/w), PGPC (1.25, 2.5, 5, 10 w/w), and MDP (1.25, 2.5, 5, 10 w/w) for 20 minutes at room temperature, followed by blue native PAGE analysis.

### Recombinant protein production and purification

pFastBac plasmid encoding full-length caspase-11 with a C254A mutation and N-terminal His-tag was a gift from Jianbin Ruan (University of Connecticut), and the protein was produced in Sf9 insect cells. Chemically competent DH10Bac cells (ThermoFisher) were transformed with the pFastBac vector encoding caspase-11(C254A) and plated on LB agar plates containing 25 µg/ml kanamycin, 10 µg/ml tetracycline, 7 µg/ml gentamycin, 50 µg/ml X-Gal, and 40 µg/ml IPTG, and incubated overnight at 37 °C for blue/white screening. A single white colony was inoculated into 20 ml of LB medium with 25 µg/ml kanamycin, 10 µg/ml tetracycline, and 7 µg/ml gentamycin and incubated overnight at 37 °C and 250 rpm. Bacmid DNA was isolated using buffer components from the GeneJET Plasmid Miniprep kit (ThermoFisher) and precipitated using isopropanol. The DNA pellet was washed once with 70% ethanol, air-dried, and resuspended in 40 µl of sterile, ultrapure water. Ten microliters of bacmid DNA-containing water were diluted with 100 µl of Hyclone SFX insect cell media. Ten microliters of CellFectin II (ThermoFisher) transfection reagent were mixed with 100 µl of media and added to the DNA mixture. After incubating at room temperature for 30 minutes, 100 µl of the transfection mix was added dropwise to 0.8 x 10⁶ Sf9 cells seeded in a 6-well plate with 3 ml of media. After 2–3 days of incubation at 28 °C, the baculovirus-containing supernatant was collected. Virus-containing supernatants from two wells were combined, filtered using a 0.45 µm syringe filter, and stored at 4 °C. To amplify the virus, Sf9 cells were grown to a density of 1.0 x 10⁶ cells/ml in 25 ml, infected with 2 ml of the initial virus, and incubated at 28 °C and 120 rpm. After 3 days, virus was collected by centrifuging the cells at 1000g for 10 minutes, and the supernatant was filtered (0.45 µm syringe filter) and stored at 4 °C. For protein expression, Sf9 cells were seeded at a density of 1.5 x 10⁶ cells/ml in a 1-liter suspension culture, infected with 1% (v/v) of the amplified virus, and incubated at 28 °C and 120 rpm. After 96 hours of incubation, cell proliferation arrested and cells were harvested by centrifugation at 1000g for 15 minutes. Cell pellets were frozen in liquid nitrogen and stored at –20 °C until protein purification. For protein purification, frozen cell pellets were resuspended in 100 ml of resuspension buffer (25 mM HEPES-NaOH pH 7.4, 150 mM NaCl, and 10 mM imidazole) and lysed by ultrasonication. The lysate was clarified by centrifugation at 20,000g for 45 minutes at 4 °C. The supernatant was incubated with 1 ml of Ni-NTA agarose beads (Qiagen) pre-equilibrated in resuspension buffer for 1 hour at 4 °C with gentle rotation. The beads were collected by centrifugation, resuspended in 10 ml of resuspension buffer, and transferred to a gravity flow column. The beads were washed with 20 bed volumes of wash buffer (25 mM HEPES-NaOH pH 7.4, 400 mM NaCl, and 25 mM imidazole), and bound protein was eluted in 10 ml of elution buffer (25 mM HEPES-NaOH pH 7.4, 150 mM NaCl, and 250 mM imidazole). Eluted protein was concentrated using Amicon Ultra-15 centrifugal filter units with a 10 kDa cut-off (EMD Millipore) and further purified by size-exclusion chromatography using a BioRad NGC Quest10 Chromatography system equipped with a Superdex 200 Increase 10/300 column in SEC buffer (25 mM HEPES-NaOH pH 7.4, 150 mM NaCl). Fractions containing pure caspase-11 were identified by SDS-PAGE and InstantBlue staining (Expedeon) and combined. Glycerol was added to the combined fractions to a final concentration of 10%. Protein concentration was measured by absorbance at 280 nm using a Nanodrop device, corrected for the protein-specific extinction coefficient. Aliquots with a protein concentration between 0.5 and 1 mg/ml were snap-frozen in liquid nitrogen and stored at –80 °C.

### Bioinformatic analysis

Predicting structure using AlphaFold: the first 90 amino acids were defined as the CARD of caspase-11. The amino acid sequence was input into the AlphaFold Colab Page (version 1.5.2) using MMseqs2 in unpaired mode. Models were ranked based on the predicted local distance difference test (plddt) scores, with auto-selection for the model type and the number of recycles. The PDB file of the top-ranked model was selected as the predicted structure for caspase-11 CARD. The predicted structure of caspase-11 CARD was used as the target for ligand docking in SwissDock (Old Version: old.swissdock.ch/docking). Sodium dodecyl sulfate (SDS, ZINC1532179) was selected as the ligand. Orthologs searching using Foldseek (https://search.foldseek.com/search): The predicted structure of caspase-11 CARD was used as the input for Foldseek server to search on AlphaFold/UniProt50, CATH50, and PDB100. The returned structures were examined manually. Candidates were cloned to pMSCV-IRES-GFP backbone. Sequence alignment was performed using CLUSTALW (https://www.genome.jp/tools-bin/clustalw), illustrated using ESPript3 (https://espript.ibcp.fr/ESPript/ESPript/).

### Limited trypsin digestion with recombinant protein and mass spectrometry

2 µg of recombinant full-length catalytically inactive (C254A) caspase-11 were reconstituted in 10 µL of LTD buffer and incubated with 10 µg of PGPC (1 mg/mL) at room temperature for 20 minutes. Five microliters of freshly prepared trypsin (50 µg/mL) were added to each reaction, followed by incubation. The reaction was stopped by adding 2X Laemmli buffer after 2 minutes and heating at 95 °C for 5 minutes. Fifteen microliters of each sample were loaded onto NuPAGE™ Bis-Tris Mini Protein Gels, 4–12% (1.0 mm, NP0321BOX) and electrophoresed. Corresponding protein bands were visualized using InstantBlue® Coomassie Protein Stain (abcam, ISB1L) and rinsed with distilled water. The corresponding gel bands were excised and sent for LC-MS/MS.

### Protein Sequence Analysis by LC-MS/MS

Excised gel bands were cut into approximately 1 mm^3^ pieces. Gel pieces were then subjected to a modified in-gel trypsin digestion procedure. Gel pieces were washed and dehydrated with acetonitrile for 10 min. followed by removal of acetonitrile. Pieces were then completely dried in a speed-vac. Rehydration of the gel pieces was with 50 mM ammonium bicarbonate solution containing 12.5 ng/µl modified sequencing-grade trypsin (Promega, Madison, WI) at 4°C. After 45 min., the excess trypsin solution was removed and replaced with 50 mM ammonium bicarbonate solution to just cover the gel pieces. Samples were then placed in a 37°C room overnight. Peptides were later extracted by removing the ammonium bicarbonate solution, followed by one wash with a solution containing 50% acetonitrile and 1% formic acid. The extracts were then dried in a speed-vac (∼1 hr). The samples were then stored at 4°C until analysis. On the day of analysis the samples were reconstituted in 5 - 10 µl of HPLC solvent A (2.5% acetonitrile, 0.1% formic acid). A nano-scale reverse-phase HPLC capillary column was created by packing 2.6 µm C18 spherical silica beads into a fused silica capillary (100 µm inner diameter x ∼30 cm length) with a flame-drawn tip. After equilibrating the column each sample was loaded via a Famos auto sampler (LC Packings, San Francisco CA) onto the column. A gradient was formed and peptides were eluted with increasing concentrations of solvent B (97.5% acetonitrile, 0.1% formic acid). As peptides eluted they were subjected to electrospray ionization and then entered into a Velos Orbitrap Pro ion-trap mass spectrometer (Thermo Fisher Scientific, Waltham, MA). Peptides were detected, isolated, and fragmented to produce a tandem mass spectrum of specific fragment ions for each peptide. Peptide sequences (and hence protein identity) were determined by matching protein databases with the acquired fragmentation pattern by the software program, Sequest (Thermo Fisher Scientific, Waltham, MA). All databases include a reversed version of all the sequences and the data was filtered to between a one and two percent peptide false discovery rate.

### Recombinant CARD and liquid chromatography

pFastBac plasmid encoding caspase-11 CARD domain was expression and purification similar as full-length caspase-11 with a C254A mutation. The purified caspase-11 CARD was further purified with size exclusion chromatography (Superdex 200(10/300)) with buffer A (150mM NaCl, 25mM Tris-HCl, pH8.0). Fractions containing pure caspase-11 were identified by SDS-PAGE. Take pure protein (final concentration 20uM) and mixture with PGPC (final concentration 400uM) according 1:20 molar ratio in a total volume of 500uL and incubate for 10min. The mixture was then sonication for 1min (10% power, 1s on, 3s off). Then the sample centrifuge for 30min, 1400rpm. The sample finally was loaded on the Superdex75(10/300) with buffer A (150mM NaCl, 25mM Tris-HCl, pH8.0). Casp11 CARD alone sample without PGPC was done with all the processing steps at the same time.

### Microscale thermophoresis

Plasmid encoding MBP-tagged Caspase11 CARD protein containing an extra cysteine amino acid was applied for Caspase11 CARD expression and purification. The pure purified protein from gel filtration mixture with Alexa Fluor™ 647 C2 Maleimide (Invitrogen, A20347) at the molar ratio of 1:2, cover the tube with aluminium-foil paper to avoid the light. Incubate the mixture at 4℃ overnight. Load the labeled mixture to Superdex 200 (10/300) with buffer B (25mM HEPES, pH7.5; 150mM NaCl) to get pure well labelled Caspase11 CARD protein for microscale thermophoresis assay. Microscale thermophoresis (MST) experiments were performed according to the manufacturer’s instructions (NanoTemper Technologies, Monolith). The protein concentration applied in the assay is 50nM and the ligand was started at highest concentration of 500uM. A serial dilution of the ligand was applied in the assay. After incubation of protein and ligands, samples were loaded filled into MST Premium coated capillaries (NanoTemper Technologies monolith, Act# MO-K025) and measurements were taken at a constant temperature of 16 °C. The MO.Affinity Analysis Software (NanoTemper) was used to analyze the binding affinity and the Kd value (dissociation constant) with the fitting model. The thermophoretic movement changes were represented by the normalized fluorescence ratio difference before and after heating (ΔFnorm [‰]) and plotted against ligand concentration. The binding curve generated was then fitted using software to determine binding constants and calculate the dissociation constant (Kd).

### Cell Culture, Retroviral Transduction, and Selection of Caspase-11-/-Macrophages

Caspase-11⁻/⁻ immortalized macrophages were cultured in Dulbecco’s Modified Eagle’s Medium (DMEM) supplemented with 10% fetal bovine serum (FBS), penicillin-streptomycin (Pen-Strep), l-glutamine, and sodium pyruvate. 293T cells were used as a packaging cell line for retroviral vectors. For the production of retroviral particles, 7.5 × 10^5^ HEK293T cells were seeded in a each well of 6-well plate. After overnight incubation at 37°C, the cells were transfected with 1.5 µg of pMSCV IRES EGFP encoding the protein of interest, 1 µg of pCL-ECO, and 0.5 µg of pCMV-VSVG using 9 µl Lipofectamine 2000 in 150 µl total volume (Thermo Fisher Scientific) according to the manufacturer’s instructions. After 16 hours at 37°C, the media were replaced with 2 ml of fresh complete media with additional FBS (30% in final concentration). The virus-containing supernatant was collected 24 hours after the media change. Supernatants were clarified by centrifugation at 400g for 5 minutes and filtered through a 0.45-µm polyvinylidene difluoride (PVDF) syringe filter. Approximately 1 × 10^5^ caspase-11⁻/⁻ immortalized bone marrow-derived macrophages (iBMDMs) were transduced twice on consecutive days in a 12-well plate by adding 1.5 ml of viral supernatant, supplemented with polybrene (1:2000; EMD Millipore) per well. The plates were centrifuged at 1250g for 1 hour at 30°C. GFP-positive (GFP⁺). After centrifuge, 1.0 mL of fresh media was added to each well. Cells were subjected to the second round of centrifuge with virus-containing supernatant after 24 hours. Cells were transferred to fresh media after 24 hours and were allowed to expanded to be sorted twice using FACSMelody cell sorter (BD Biosciences) to generate cell lines with stable and homogeneous expression of the target protein. Transgene expression was confirmed by immunoblotting using a rat anti-caspase-11 antibody (Cell Signaling Technologies, 14340).

### Cytoplasmic Delivery of Bacterial LPS Using the Neon Transfection System

Murine iBMDMs were transfected with bacterial LPS using the Neon Transfection System (Thermo Fisher Scientific). Cells were first primed with R848 for 3 hours, then lifted with PBS + EDTA and resuspended in R buffer at a concentration of 10 × 10⁶ cells/ml. The cell suspension was mixed with either LPS (1 µg per 1 × 10⁶ cells) or sterile PBS (1 µl per 1 × 10⁶ cells as a control). The mixture was aspirated into a 100-µl electroporation pipette tip, and electroporation was performed with two pulses, each 10 ms long, at 1400 V (unless otherwise specified). The transfected cells were transferred into media at 1 × 10⁶ cells/ml and seeded into 96-well plates at 100 µl per well. For the nigericin treated control, lifted iBMDM was added directly to nigericin contraining media (10 μM) at the same cell concentration as the LPS electroporated cells. Cell death and IL-1β release were measured 6 hours later using an LDH assay and a Lumit assay, respectively.

### Cell death assays, IL-1β release, and trypsin activity quantification

Cell lysis was assessed using the CyQuant LDH Cytotoxicity Assay Kit (Thermo Fisher Scientific). 25 µl microliters of supernatant were mixed with 25 µl of LDH assay buffer and incubated at room temperature for 15 minutes. Absorbance was measured at 490 and 680 nm using a Tecan Spark plate reader, and the signal was normalized to the lysis controls. IL-1β levels were quantified using the Promega Lumit™ Mouse IL-1β Immunoassay. 25 µl of sample were mixed with 25 µl of antibody mixture and incubated at 37°C for 1 hour. T12.5 µl of luminescence substrate were added, and the luminescent signal was measured using a Tecan Spark plate reader. Trypsin activity was measured using trypsin activity assay kit (ab102531).

### Statistical Analysis

Statistical significance was assessed using one-way ANOVA followed by Tukey’s multiple comparison test. A P-value of less than 0.05 was considered statistically significant. All statistical analyses were performed using GraphPad Prism data analysis software. Data are representative of at least two independent experiments, and results with error bars are presented as means ± SEM.

## ACKNOWLEDGMENTS

We thank members of the Kagan lab for helpful discussions. This study was supported by NIH grants AI167993, AI116550, and DK34854 to J.C.K.

---

**Supplementary Figure 1:**
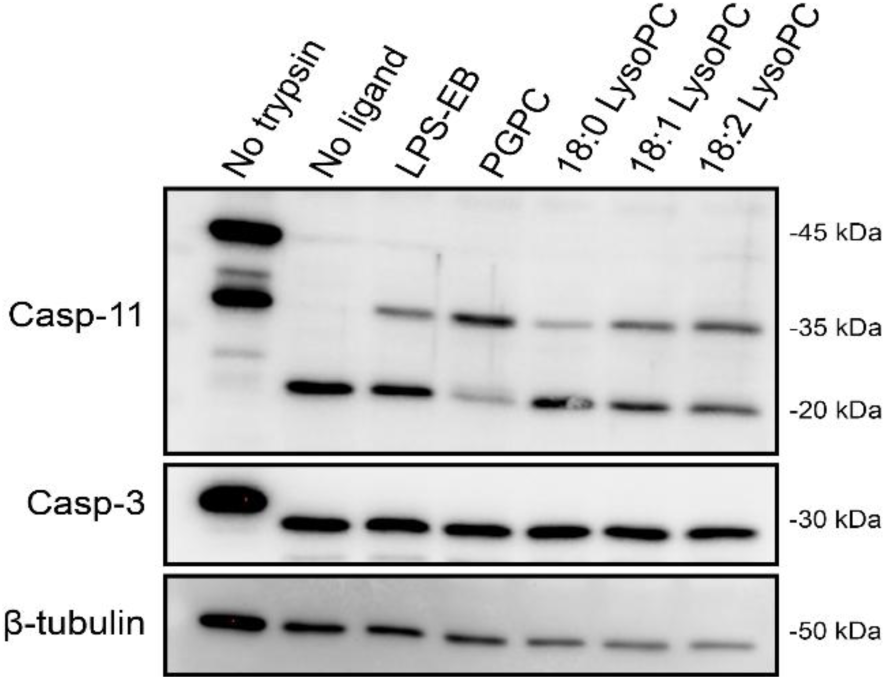
Unsaturation degree is not a determinant of caspase-11 binding. (A) 293T cell lysates expressing C254A caspase-11 were incubated with LPS (10 µg/mL) and PGPC, 18:0 LysoPC, 18:1 LysoPC, 18:2 LysoPC (100 µg/mL) and were subjected to limited trypsin digestion followed by immune blot to detect caspase-11.

**Supplementary Figure 2:**
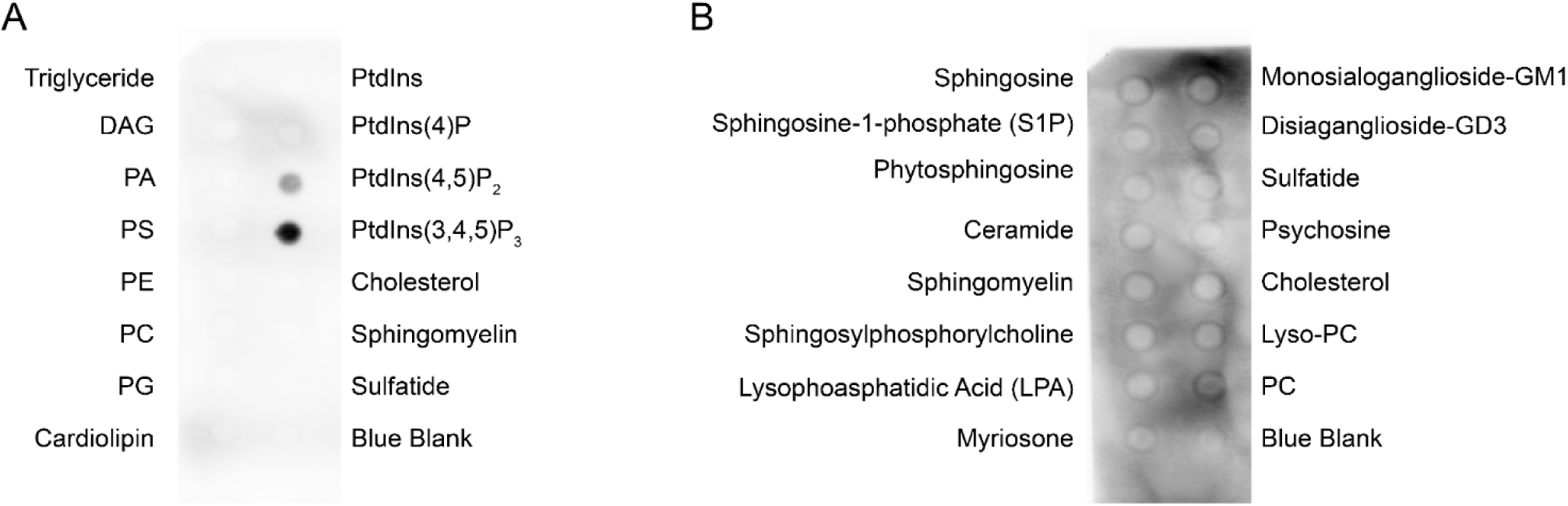
Recombinant caspase-11 binds to phosphorylated self-lipids. (A) Recombinant caspase-11 was incubated with membrane lipid-strip containing various lipids found in cell membranes, followed by immunoblot detection to assess phosphoinositide binding by caspase-11. (B) Recombinant caspase-11 was incubated with membrane lipid-strip containing various lipids containing a backbone of sphingoid bases, followed by immunoblot detection to assess phosphoinositide binding by caspase-11.

